# Eco-evolutionary drivers of avian migratory connectivity

**DOI:** 10.1101/2022.07.04.497586

**Authors:** Niccolò Fattorini, Alessandra Costanzo, Andrea Romano, Diego Rubolini, Stephen Baillie, Franz Bairlein, Fernando Spina, Roberto Ambrosini

## Abstract

Understanding how individuals redistribute after migration assists in the conservation and management of mobile species, yet the eco-evolutionary drivers of migratory connectivity remain unclear. Taking advantage of an exceptionally large (~150,000 individuals, 83 species) and more-than-a-century long dataset of bird ringing encounters, we investigated determinants of avian migratory connectivity on both short- and long-distance migrants. Most species exhibited significant connectivity likely due to the large intraspecific variability in migration strategies, which often led to the identification of distinct migratory populations. Migratory connectivity was strongly predicted by geography, especially migration distance, but it was evolutionary labile and weakly influenced by biological traits due to flexibility in migratory behaviour. By unravelling determinants of migratory connectivity we improve knowledge about the resilience of migrants to ecological perturbations. Also, our population-level analysis provides a critical tool to inform transboundary conservation and management strategies by explicitly considering the large intraspecific variability of avian migration.

## Introduction

Animal migration is a widespread phenomenon which has evolved as an adaptive response to spatiotemporal variations in resources (Dingle & Drake 2007). Ongoing environmental changes are disrupting migration strategies and threatening migratory populations at the global scale, with dramatic declines in the abundance of migratory species and a reduction of the occurrence of migratory behaviours, leading to an increasing number of populations of formerly migratory species becoming resident (Wilcove & Wikelski 2008; Robinson et al. 2009; Visser et al. 2009; Pulido and Berthold 2010; Runge et al. 2014; Morganti 2015; Shaw 2016; Buchan et al. 2020). Migratory connectivity, i.e. the spatiotemporal linkage between breeding and nonbreeding ranges where individuals spend different phases of their annual cycle (Webster et al. 2009), is an important concept offering the possibility to assist in the conservation of migratory species ranging from invertebrates (Gao et al. 2020) to marine mammals (Dunn et al. 2019). This ecological property quantifies the degree of population mixing after migration and has become critical to understand how ecological perturbations in the nonbreeding range may affect the fitness and population dynamics of migrants (Taylor & Norris 2010). Particularly among birds, strongly connected migratory populations have been suggested to be more vulnerable to environmental changes because any change in the nonbreeding grounds would affect the entire population (Briedis & Bauer 2018; Ambrosini et al. 2019). In contrast, the negative consequences of any environmental alteration may be buffered in loosely connected populations because only part of a breeding population would experience such changes (Briedis & Bauer 2018; Ambrosini et al. 2019). Hence, understanding the processes affecting the population dynamics of migratory species requires an improved knowledge of the ecological and evolutionary causes of migratory connectivity (Boulet & Norris 2006; Patchett et al. 2018; Beresford et al. 2019).

Although previous studies have identified some geographical and bio-energetic conditions that affect the strength of avian migratory connectivity (Finch et al. 2018; Somveille et al. 2021), the underlying eco-evolutionary drivers are poorly understood. In particular, no interspecific comparative analysis has been conducted, though such analysis would be essential to disentangle the potential evolutionary and ecological drivers of the extent of migratory connectivity in different species. In fact, both temporal and spatial migration patterns are primarily inherited in birds (Liedvogel et al. 2011; Åkesson & Helm 2020; Gu et al. 2021), leading to hypothesize the existence of a phylogenetic signal on the strength of migratory connectivity. Additionally, since avian migration is largely influenced by life-history traits such as niche specialisation (Reif et al. 2016) and body mass-dependent energetic costs of aerial locomotion (Hein et al. 2012), these and other similar ecological factors could play critical roles in determining how birds redistribute between breeding and nonbreeding ranges.

Here, we took advantage of over a century-long collection of ringing encounters of European breeding birds (du Feu et al. 2016) to investigate the eco-evolutionary determinants of avian migratory connectivity. We worked at the scale of geographical populations along a gradient of migratory behaviours ranging from short-to long-distance migrants, as well as from partial to fully migratory species, and tested a set of hypotheses on the factors potentially explaining the strength of migratory connectivity. We simultaneously tested for phylogenetic, geographic and life-history effects using a mixed model approach based on a larger number of species, populations and individuals compared to previous studies.

Despite available theories well predict the observed patterns of avian migratory connectivity strength at continental scales (Finch et al. 2018; Somveille et al. 2021), multi-species information based on empirical data collected within the European-African migration system is scarce. Over the last decades, European populations of migratory birds, especially long-distance ones, have declined substantially, likely due to habitat loss or deterioration in the nonbreeding grounds (Sanderson et al. 2006; Vickery et al. 2014; Beresford et al. 2019; Howard et al. 2020). Migratory connectivity seems to play a key-role in such declines (Patchett et al. 2018), and previous studies call for gaining knowledge on how birds mix between breeding and nonbreeding grounds to assist the conservation of European migrants (Beresford et al. 2019). Furthermore, long-distance migrants in the European-African system seem to have already responded to past climatic perturbations occurring in Africa through the evolution of low migratory connectivity as a bethedging adaptation (Cresswell 2014; Patchett et al. 2018), which makes this an ideal study system to investigate the ecological drivers of migratory connectivity. We thus focused on this migration system and used data concerning ~150,000 individual birds from 83 species belonging to 32 avian families. To the best of our knowledge, the present study involves the largest dataset ever used to investigate migratory connectivity in the animal kingdom.

Our expectations on the driving forces potentially influencing avian migratory connectivity originate from evidence provided by earlier research, but also extend towards unexplored effects of life-history traits stemming from ecological theory, as summarized in Table 1 (see also Appendix S1, Supporting Information, for details). By unravelling eco-evolutionary determinants of avian migratory connectivity, the present study improves our understanding about the resilience of migratory birds to ecological perturbations, and provides a critical tool to inform transboundary conservation and management strategies.

**Table 1.**
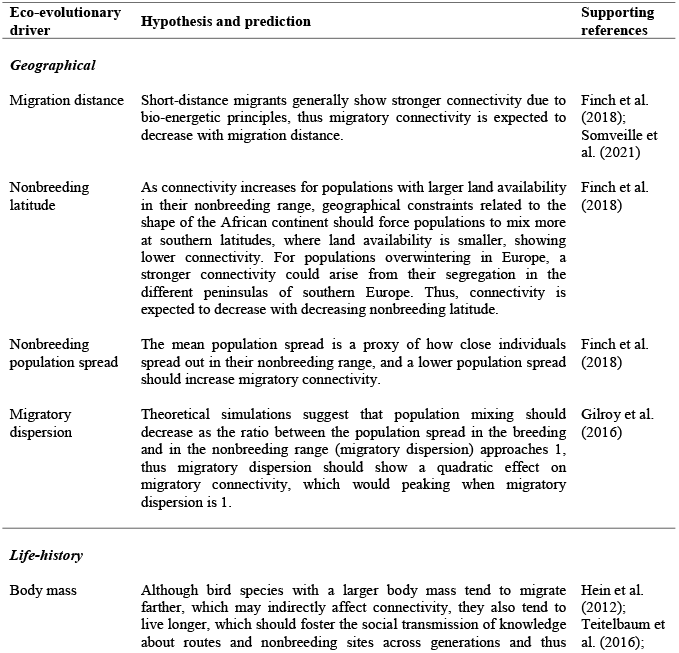
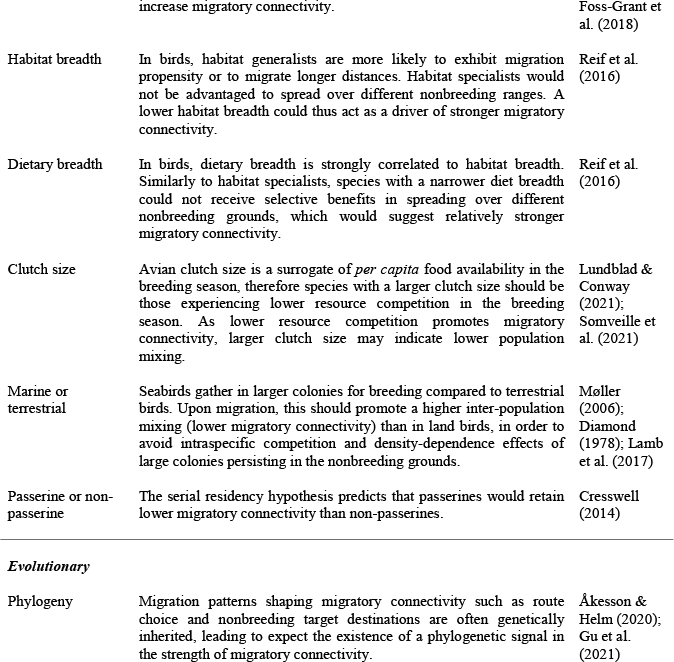
Hypotheses tested about the effects of potential eco-evolutionary drivers of avian migratory connectivity. See also Appendix S1, for an extended explanation.

## Materials and Methods

### Migratory connectivity analysis

Initially, we assessed the strength of migratory connectivity for 137 bird species filtering more than 12 million ringing encounters obtained from the EURING databank (du Feu et al. 2016; https://euring.org) and spanning over one century (1900 2019). Ringing data are largely heterogeneous as to individual encounter condition and circumstances, such as those of birds in poor health status or kept in captivity or manipulated for long, or those for which the place/time of recovery was not determined accurately, therefore preliminary data filtering is necessary to reduce heterogeneity (e.g. Paradis et al. 1998). Moreover, ringing encounters collected outside species-specific breeding or nonbreeding grounds and periods, potentially resulting from occasional movements, would introduce spatial and temporal biases in migratory connectivity analysis (e.g. Somveille et al. 2021). Hence, we implemented a robust data selection procedure to discard ringing encounters potentially affecting our estimates of migratory connectivity (Appendix S2). Our conservative approach relied on 21 condition-based selection criteria and applied a spatiotemporal masking for each species, discarding on average 97.2% of ringing encounters per species (range: 58% □ 99%; see Ambrosini et al. 2016 for a similar data selection procedure). Selected data included 371,090 individuals from 137 species (range: 20 □ 36,506 individuals/species; mean ± SE: 2,708 ± 480 individuals/species) encountered between 1917 2019.

For each species, we assessed the strength of migratory connectivity according to Ambrosini et al. (2009). The method was developed using ringing encounters and has been used consistently to estimate migratory connectivity at various geographical scales, and for both single- and multi-species analyses (e.g. Ambrosini et al. 2016; Finch et al. 2018; Sarà et al. 2019; Knight et al. 2021; Somveille et al. 2021). The strength of migratory connectivity was quantified through the Mantel correlation coefficient (*r_M_*), and the probability of a positive connectivity was tested by a one-tailed permutation test (Ambrosini et al. 2009) because negative values in the strength of migratory connectivity are not biologically meaningful (Cohen et al. 2018). For those species showing significant connectivity, eight k-mean cluster analyses (pre-defined number of clusters: 2□9) were performed to identify clusters of individuals that tend to gather in separate groups in the breeding and nonbreeding ranges (Ambrosini et al. 2009). The best clustering structure was identified as that maximising the overall average silhouette width (*oasw;* Rousseeuw 1987), a metric showing the best performance amongst clustering validity indices (Arbelaitz et al. 2013). We stress that a reasonable or strong clustering structure (*oasw* ≥ 0.5) suggests the presence of geographical populations that differ in their breeding or nonbreeding grounds. Thus, for species showing distinct clusters, we re-calculated *r_M_* on each cluster separately and we used these values rather than that calculated on the whole species in the subsequent analyses. The rationale behind this choice is that migration strategies can largely differ among geographical populations of the same species, so the analyses to assess the ecological drivers of migratory connectivity should be based on either the *r_M_* value calculated on each geographical population, for those species showing distinct clusters (Fig. 1a,b), or on the *r_M_* value calculated on all the encounters, for those species that were not spatially clustered into distinct populations (Fig. 1c,d). For the latter species (*oasw* < 0.5), we thus considered all individuals as belonging to a single cluster, independently of the significance of *r_M_*. Hereafter we will refer to a ‘geographical population’ as either (1) the ensemble of all the ring encounters of a species, if the species showed non-significant connectivity or a weak clustering structure by which all individuals can be considered to belong to the same geographical population, or (2) each of the clusters identified by the migratory connectivity analysis at the species level if the species showed a significant connectivity and a reasonable or strong clustering structure. A bootstrap procedure was also used to estimate the 95% CI of *r_M_* values.

**Figure 1.**
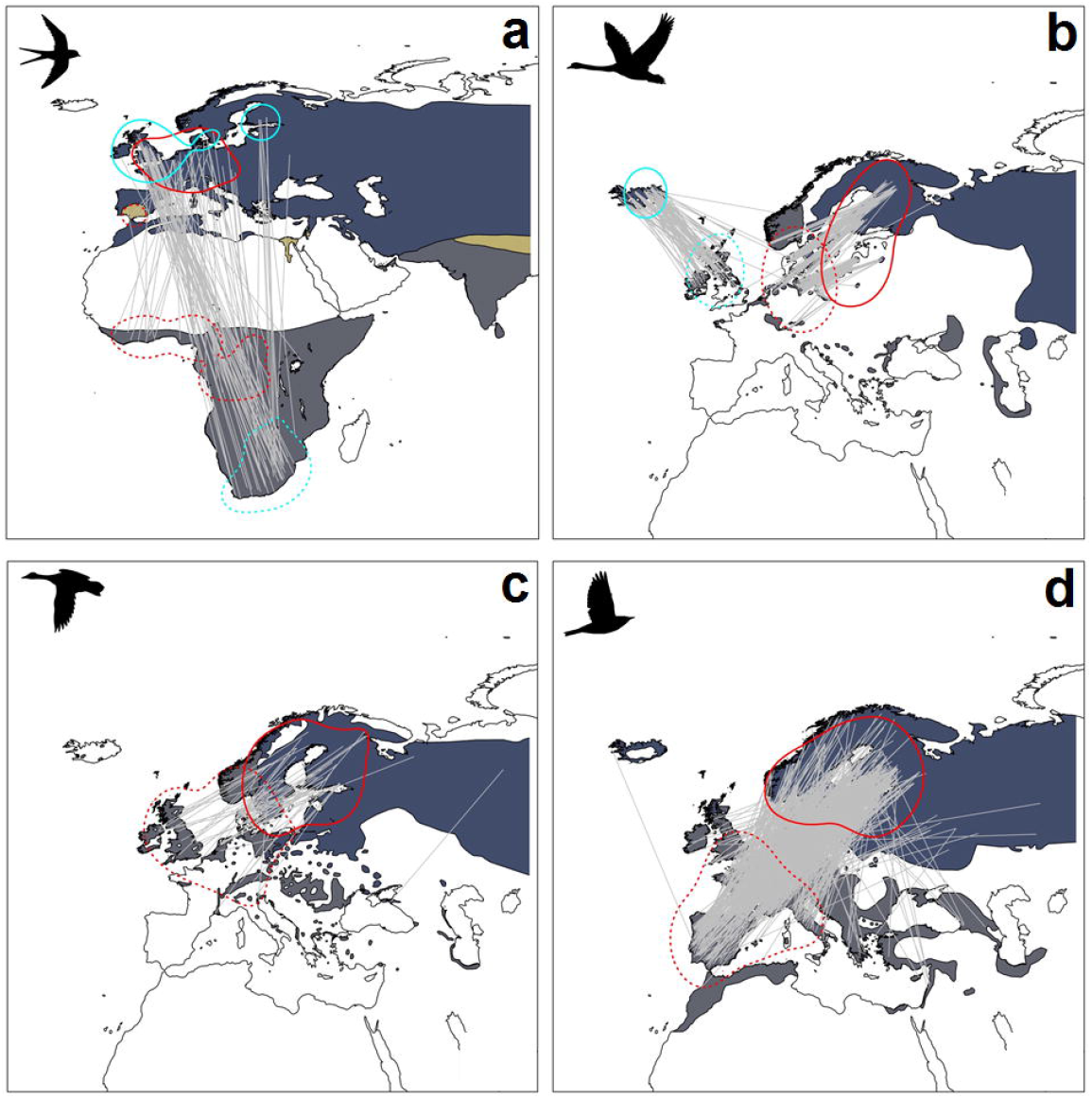
Migratory connectivity maps of representative bird species showing (a) weak connectivity and strong clustering (*Hirundo rustica: r_M_* = 0.15, P = 0.004, n = 96, *oasw* = 0.61); (b) strong connectivity and strong clustering (*Cygnus cygnus: r_M_* = 0.87, P < 0.001, n = 509, *oasw* = 0.76); (c) moderate connectivity and weak clustering (*Bucephala clangula: r_M_* = 0.40, P = 0.003, n = 69, *oasw* = 0.34); (d) no connectivity (*Turdus iliacus: r_M_* = 0.03, P = 0.13, n = 683). The grey lines connect individual breeding sites and nonbreeding destinations, while differently coloured kernel contours encompass 95% ring encounters of different geographical populations (solid contour: breeding, dotted contour: nonbreeding). The species distribution range is also shown (breeding range: blue; non-breeding range: dark grey; resident range: beige; from BirdLife International, 2019). Connectivity maps for all species can be accessed at https://migrationatlas.org/ (Spina et al. 2022).

We classified the geographical populations identified by the migratory connectivity analysis as migratory or resident based on the extent of the overlap of their breeding and nonbreeding ranges (see Appendix S3, for details). As we were interested in shedding light on the factors influencing the strength of the connectivity of migratory species, being them full or partial migrants, we discarded the species with only resident geographical populations in our dataset. The analyses were therefore run including all the geographical populations of those species that had at least one geographical population classified as migratory. Additionally, we only retained geographical populations having ≥ 30 individuals re-encountered, because lower sample sizes may not provide robust connectivity estimates (Ambrosini et al. 2009). Our final dataset included 150,909 individuals from 191 geographical populations belonging to 83 species (range: 30 – 27,479 individuals/species and 1-9 geographical populations/species; mean ± SE: 1,818 ± 429 individuals/species and 2.30 ± 0.21 populations/species; Appendix S4). In a second step, to assess potential differences between migratory and sedentary populations, we re-ran the analyses by removing geographical populations that were classified as resident, retaining 120,377 individuals and 150 geographical populations from 83 species.

### Phylogenetic comparative meta-analysis

A phylogenetic meta-analysis was conducted using the *metafor* R package (Viechtbauer 2010). Because *r_M_* values are correlation coefficients, they were transformed into *Z_r_* using Fisher transformation. We then fitted a phylogenetic mixed model where the variance components of the random part allow calculating how much variance is attributable to the phylogeny (phylogenetic hereditability, *H^2^*) while considering multiple *Z_r_* values for the same species and accounting for the fixed effects included in the model (*H^2^* is equivalent to Pagel’s λ; see Nakagawa & Santos 2012). *H^2^* was calculated as the ratio between the variance due to phylogeny and all the variance components in the model (Nakagawa & Santos 2012). To account for phylogeny, we built a 50% majority rule-consensus tree using 10,000 phylogenetic trees (Hackett et al. 2008) retrieved from www.birdtree.org, as recommended for avian comparative studies (Rubolini et al. 2015).

In a second model, we also included as fixed effects a set of moderators that may influence migratory connectivity according to our hypotheses stemming from ecological theory (Table 1 and Appendix S1). For each geographical population, the mean (orthodromic) migration distance, the population spread (mean interindividual pairwise distance in the nonbreeding population range, following Finch et al. 2018), the mean nonbreeding latitude, and both the linear and quadratic effects of migratory dispersion (see Table 1; dispersion was computed as the ratio between breeding and nonbreeding population spread, both calculated as above) were obtained from the position of the ringing encounters used in the analyses. Life-history traits (i.e. body mass, clutch size, habitat and dietary breadths) were compiled from the literature (Appendix S5). Finally, we included two dichotomous moderators indicating whether a species was marine or non-marine, and whether it was a passerine or a non-passerine species. Preliminary, we explored the relationships between *Z_r_* values and predictors, and applied logarithmic data transformation whenever we detected nonlinear effects (i.e., to migration distance and nonbreeding population spread; Appendix S6-S7). All predictors (including binary ones) were scaled, and we found no multicollinearity between predictors (Pearson’s |r| ≤ 0.55; VIF ≤ 2.7). Preliminary exploration also suggested an heterogeneity of variance in *Z_r_* values between marine and non-marine species and between passerines and non-passerines (Appendix S7). Hence, we allowed for heterogeneity of variance between these groups by entering the binary predictors as inner variables in the random part of the model and setting a diagonal covariance structure for the variance-covariance matrix. This allowed the model estimating different variances for each level of these predictors. We also scaled *Z_r_* values by the inverse of their variance (equal to *N* □ 3, where *N* is the number of individuals in a geographical population). Degrees of freedom were calculated with the containment method that offers a better control of the Type I error rate and produces confidence intervals with closer-to-nominal coverage rates (https://wviechtb.github.io/metafor/reference/index.html). Model were fitted using REML, and t-values were used as measures of effect size.

## Results

### Model without moderators

Overall, migratory connectivity was moderate and significantly larger than zero (estimated *Z_r_* = 0.471 ± 0.119 SE, *t_82_* = 3.965, P < 0.001, corresponding to *r_M_* = 0.439, 95% CI: 0.232 – 0.608; Fig. 2). Hereditability (*H^2^* = 0.204), albeit weak, was significant (Likelihood Ratio Test with a model not accounting for species phylogeny, *χ*^2^ = 8.077, *df* = 1, P = 0.004), indicating the presence of a phylogenetic signal in the strength of migratory connectivity.

**Figure 2.**
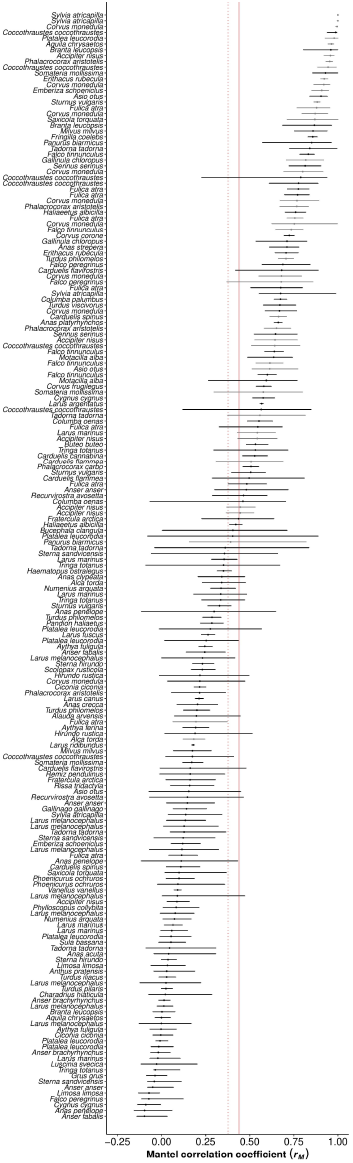
Forest plot showing the strength of migratory connectivity (*r_M_*) estimated for 191 geographical populations of 83 bird species, with error bars depicting bootstrapped 95% CIs of *r_M_* values (black: migratory populations; grey: sedentary populations). Values are sorted according to connectivity strength. Red lines show the mean connectivity strength estimated by the null models fitted while considering migratory and resident populations (solid) or only migratory populations (dotted).

### Model with moderators

We found a negative effect of migration distance and a positive effect of population spread on the strength of migratory connectivity, with migration distance having more than a threefold stronger effect than population spread (Table 2; Fig. 3). Additionally, there was a significant quadratic effect of the ratio between breeding and nonbreeding population spread (migratory dispersion) though this moderator had a lower effect size than that of migration distance (Table 2; Fig. 3). Migratory connectivity also increased with decreasing nonbreeding latitude (Table 2; Fig. 3), implying that populations with nonbreeding ranges at southern latitudes showed a lower population mixing. Despite the effect of nonbreeding latitude was the weakest amongst geographic predictors, this result did not change when we excluded geographical populations with nonbreeding latitude < 30° N from the analysis (details not shown). As to species-specific biological traits, only habitat breadth was significantly related to migratory connectivity, whereby habitat generalists showed a stronger migratory connectivity (Table 2; Fig. 3). Similarly to the null model, migratory connectivity was generally positive and significantly larger than zero across all populations, suggesting that individuals tend to maintain their reciprocal positions within the geographical populations (Table 2; since all our predictors were scaled before the analysis, the intercept reflects the general mean value across all clusters). Moreover, hereditability was strong and significant (*H^2^* = 0.642, χ^2^ = 4.261, *df* = 1, P = 0.039). Overall, model performance was high (observed *vs* predicted values: *R^2^* = 0.80; Appendix S7) and model diagnostics indicated the robustness of our meta-analysis (Appendix S9). However, the test for residual heterogeneity was significant (QE = 3772.8032, *df* = 179, P < 0.001), possibly indicating that unaccounted moderators also influenced the strength of migratory connectivity.

**Figure 3.**
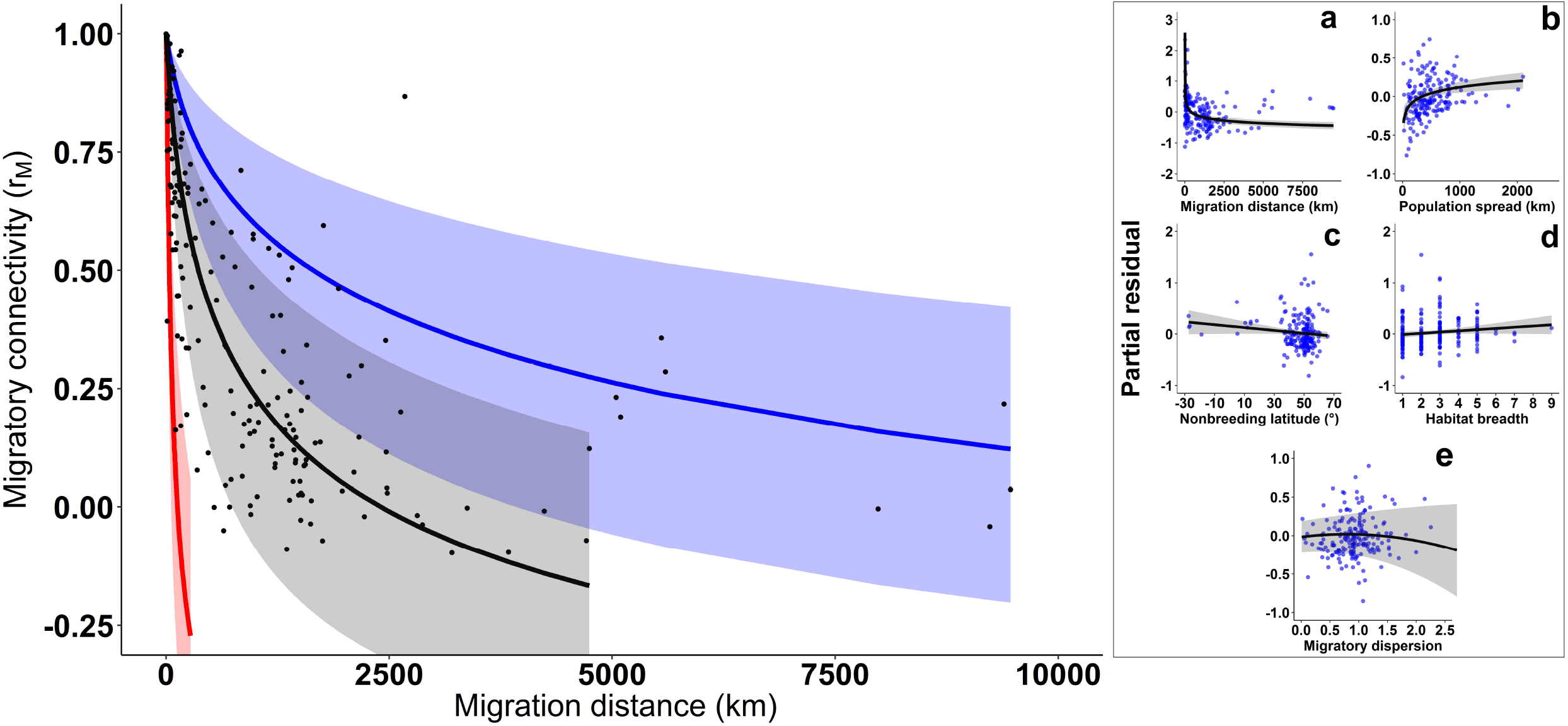
The strength of migratory connectivity (*r_M_*) predicted in relation to the covariates with the strongest effect sizes, i.e., orthodromic migration distance and nonbreeding population spread. The latter is categorized into levels corresponding to the minimum (red), median (grey), and maximum (blue) observed values: 16.02 km, 423.99 km, and 2100.41 km. The inset (a-e) shows the net effects of covariates significantly influencing the extent of migratory connectivity, as partial residuals. For modelling, Fisher Z-transformation was applied to *r_M_* values, and logarithmic transformations were applied to migration distance and migratory dispersion (see Appendix S6-S7), whilst figure and inset depict back-transformed predictions. Lines and bands: predicted values and 95% confidence intervals. Dots: observed values.

When we excluded sedentary populations from the analysis, the effect of migratory dispersion disappeared, while the effects of other geographical predictors did not change (Appendix S8). Moreover, we did not find significant effects of life-history traits and phylogeny (Appendix S8) in influencing migratory connectivity.

**Table 1.**
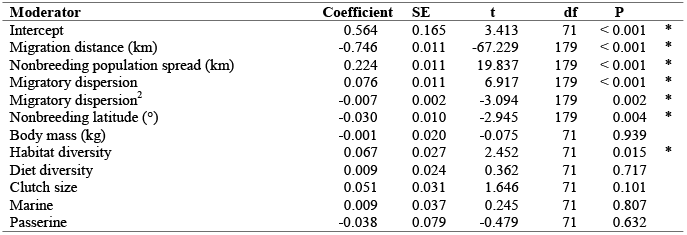
Parameters estimated from the phylogenetic meta-analytic mixed model explaining interspecific variation in the strength of migratory connectivity (as Fisher Z-transformation of *r_M_* value) across 191 geographical populations belonging to 83 bird species. Transformations were applied to ‘Migration distance’ and ‘Nonbreeding population spread’ (see Appendix S6-S7), while the second order polynomial term of ‘Migratory dispersion’ was included to account for quadratic effects (Table 1). All moderators are scaled. An asterisk marks significant (P < 0.05) moderators.

## Discussion

### The key role of geography in shaping migratory connectivity

Finch et al. (2018) have shown that low migratory connectivity is common in long-distance migrants, which spread out less extensively in the nonbreeding grounds due to reduced land availability as approaching the southern portion of the African continent. Our work generalized this result also to short-distance migrants wintering in southern Europe or Mediterranean Africa, showing that migratory connectivity was stronger in the geographical populations that spread out more extensively in the nonbreeding period, regardless of their wintering latitude and migration distance, thus supporting the hypothesis that reduced nonbreeding land availability promotes the chance of mixing. Indeed, since we estimated migratory connectivity at the geographical population level, our measure of nonbreeding population spread reflects the constraints that limit individuals to spread out solely within clusters, thus corresponding to the total nonbreeding range spread considered by Finch et al. (2018). Interestingly, when we considered both resident and migratory populations, migratory connectivity also depended on the relative population spread in the breeding period (i.e. migratory dispersion; Gilroy et al. 2016), whereby populations with similar extents of spread in seasonal ranges tended to mix less. However this effect did not occur when only migratory populations were considered, suggesting that population mixing may occur regardless of the opportunity to spread in the nonbreeding grounds relative to the breeding ones.

Previous studies have considered nonbreeding latitude, nonbreeding land availability and migration distance as a joint driving force in shaping migratory connectivity because, in the European-African migration system, land availability typically decreases at southern latitudes and long-distance migrants are those spending the nonbreeding period at the southernmost sites (Finch et al. 2018). By examining populations with nonbreeding latitudes spanning from Central Europe to Southern Africa, we disentangled the relative contribution of these factors. Contrary to our initial hypotheses, we found opposite effects of nonbreeding population spread and latitude on the strength of migratory connectivity. Bird populations wintering from Northern Europe to South Africa thus seem to show increasing migratory connectivity independently of population spread and migration distance. Additionally, our analysis quantified the nonlinear, negative effect of migration distance on the strength of migratory connectivity, which was the geographical driver with the largest effect size. Birds seem to minimize energetic costs of seasonal relocation following an optimal redistribution model, implying that individuals tend to migrate along the shortest available path due to the increasing energy expenditure with longer migration distances (Somveille et al. 2021). Our data showed that the strength of migratory connectivity drops almost linearly up to a threshold migration distance of 2000 □ 2500 km (approximately corresponding to the length of a direct crossing of the Sahara desert), then such decrease flattens up to the maximum observed migration distance of ~9500 km. Thus, the nonlinear decrease in migratory connectivity with increasing migration distance may arise from the combined costs of avian aerial locomotion and the availability of suitable habitats for refuelling that may determine a nonlinear increase of migration costs with the distance travelled by birds (Hein 2012). The same effects may also explain, at least partly, the stronger migratory connectivity at lower latitudes: birds may be forced to cross a large ecological barrier like the Sahara desert along straight longitudinal paths, and therefore tend to maintain in the sub-Saharan grounds the reciprocal positions they have before crossing it (e.g. Sarà et al. 2019). Alternatively, geographical populations migrating to southern latitudes may retain the reciprocal distances among individuals by exploiting a larger latitudinal rather than longitudinal range, as it seems to happen for example in the barn swallow *Hirundo rustica* (Ambrosini et al. 2009).

Clearly, unaccounted predictors associated to geography may also help explain the residual heterogeneity shown by our model. As suggested by Somveille et al. (2021), avian redistribution patterns in seasonal ranges could be affected by *en route* environmental conditions experienced by migrants such as wind (Kranstauber et al. 2015; Norevik et al. 2020) and, especially in these cases, optimal migration routes may depart substantially from the shortest path connecting seasonal grounds. Indeed Kranstauber et al. (2015) showed that favourable air currents influence migratory trajectories and suggested that birds could adjust migration routes at the population level by tracking efficiently the wind-optimised route. Additionally, terrestrial birds can surf the so called ‘green wave’, following the spatiotemporal gradient of vegetation productivity while migrating or when at their nonbreeding grounds (Trierweiler et al. 2013; Kölzsch et al. 2015). These opportunities may have contributed to influence the true distance travelled by some of our investigated species and populations and, in turn, their migratory connectivity, partly incorporating the unexplained variability of our data. Although the great-circle distance between breeding and nonbreeding locations is a good proxy for the actual distance travelled by avian migrants (Hein 2012, Finch et al. 2018, Somveille et al. 2021), future works based on tracking data would be necessary to refine the investigation of the migration distance effect by considering the true distance travelled by birds. In spite of the above limitations, we should note that our empirical model showed a considerable performance, providing a reliable synthesis on the determinants of avian migratory connectivity of European-African migrants.

### Life-history and evolutionary drivers of migratory connectivity vary with migration propensity

Habitat specialists showed a lower connectivity than habitat generalists, whereas other life-history traits did not seem to play a role in shaping migratory connectivity. Bird species with a narrower habitat breadth could be constrained to concentrate in relatively small nonbreeding areas where individuals are more likely to mix. This result may also reflect an adaptive response of habitat specialists to the temporal shifts in the geographical position of suitable nonbreeding habitats that have occurred in the past due to the large variability in climate conditions, particularly rainfalls, that naturally occurred in sub-Saharan Africa. As suggested by Finch et al. (2018), under largely variable conditions, a lower migratory connectivity may promote the chance of survival of a population because only a part will suffer the negative effects of an unpredicted drought in a part of the nonbreeding ranges.

The analysis conducted on both migratory and resident populations also showed a strong phylogenetic signal in migratory connectivity. Nevertheless, when we considered only migratory populations, both the life-history traits and the phylogenetic relatedness between species were not supported as influential drivers of avian migratory connectivity likely because the strong effects of geographical variables exceeded their ones. In any case, the lack of a phylogenetic signal suggests that migratory connectivity is not a trait shared between common ancestor lineages for migratory populations, implying that the way birds redistribute during migration may be an evolutionarily labile trait. Avian migration patterns such as the time of departure from seasonal sites, route choice and nonbreeding target destinations may indeed not only be inherited genetically, but also be highly flexible (Winkler et al. 2017; Åkesson & Helm 2020). For example, they may depend on individual-specific learning capacity and social transmission within groups (Mueller et al. 2013; Teitelbaum et al. 2016; Foss-Grant et al. 2018), and change across generations with varying climatic conditions experienced *en route* or in seasonal grounds (Saino & Ambrosini 2008; Clausen et al. 2018; Lameris et al. 2018; Jiguet et al. 2919; Gu et al. 2021; Dufour et al. 2021). Migratory connectivity is an emergent ecological property ultimately determined by plastic adaptations, possibly explaining why the extent of population mixing during migration does not show phylogenetic conservatism for migratory populations. Indeed, episodes of migration loss and returns to resident behaviour have often occurred across avian migratory lineages, following adaptations to novel ecological opportunities (Dufour et al. 2020). In contrast, a more phylogenetically-predictable pattern of population mixing appeared in the analysis that included resident populations, suggesting that phylogenetically-close species may have also shared similar phylogeographic processes that have promoted their resident behaviour (Winger et al. 2019).

There are, inevitably, limitations in any comparative analysis aiming at identifying the eco-evolutionary drivers of complex ecological properties such as migratory connectivity. Although we examined a larger number of species and populations compared to previous studies, our model might be improved by considering biological traits and phylogeny at the population level. This could help assess whether migratory connectivity of closely-related geographical populations is more similar than that expected by chance, and whether it is associated to population-specific rather than species-specific characteristics. Unfortunately, avian biological traits and phylogeny at the population level are currently unavailable. Phylogeny size (the number of branches in the phylogenetic tree) may especially influence the power of phylogenetic meta-analyses (Chamberlain et al. 2012), and the detection of evolutionary processes may also show phylogenetic-scale dependence (Graham et al. 2018). Therefore, the increasing availability of genomic data may represent a future challenge to build avian phylogenies at the population level for the existing bird species. This opportunity would potentially allow performing analyses at a finer phylogenetic grain, helping to draw more robust conclusions about the phylogenetic conservatism of migratory connectivity. However, we are still unaware of any ecological study implementing phylogenetic trees at the population level, and the phylogenetic tree incorporated into our model clearly represents an advancement compared to previous research on migratory connectivity, which did not consider phylogeny.

### Conservation and management implications

Previous works investigating migratory connectivity in multiple avian species generally found moderate-to-strong connectivity at the species level (Finch et al. 2018: mean *r_M_* = ~0.3, N = 28 long-distance migrant species; Somveille et al. 2021: mean *r_M_* = ~0.7, N = 25 medium-to-long distance migrant species). Our comparative analysis was the first to be conducted on both short- and long-distance migrants and suggests that individuals tend, on average, to maintain their reciprocal positions even within geographical populations. The large intraspecific variability in avian migration strategies, through which most of our species (58%; N = 83) geographically split into distinct migratory populations, is likely to underpin this effect. By deepening the concept of migratory connectivity through an analysis able to identify migratory clusters within species, we therefore suggest that conservation and management strategies must consider this large variability occurring between populations. Gaining accurate information on migratory connectivity at the population-scale would improve the conservation of mobile species, not only because efforts can be directed toward distinct population-specific nonbreeding areas where each population spreads (Trierweiler et al. 2014; Finch et al. 2018), but also because the comprehensive knowledge of the spatial connections between and within populations would allow calibrate efforts by accounting for the dependencies among seasonal ranges (Taylor & Norris 2010; Runge et al. 2014). Similarly, estimates of migratory connectivity at the population level would be critical to inform management of migratory species e.g. by providing essential information on the spread of avian-borne diseases (Chen et al. 2005) or in refining forecasting systems of bird collisions (Van Doren & Horton 2018), thus assisting in human health and safety. Knowledge on population-level migratory connectivity, combined with data on population dynamics, would also allow to manage overabundant bird populations more effectively because coordinated and population-specific efforts could be targeted on both seasonal ranges. Thus, for bird species that segregate into distinct migratory populations during seasonal migration, improved management actions could be developed by delineating discrete, including transboundary and potentially overlapping management units (e.g. Madsen et al. 2014; Bacon et al. 2019) emerging from migratory connectivity analyses.

### Conclusions and future perspectives

Taking advantage of an exceptionally large dataset spanning a diversified assembly of migratory bird species, our analysis disentangled the underlying causes of avian migratory connectivity, suggesting that such ecological property is evolutionary labile for strictly migrant species, being rather conditional on highly variable, population-specific strategies of bird migration. Generally, our findings confirm that migratory connectivity is explained chiefly by geographic forces which, in turn, are proxies for the energetic costs that individuals face in relocating between seasonal ranges (Somveille et al. 2021) and for land availability in nonbreeding grounds (Finch et al. 2018). For the first time, our approach sheds light on the relative contribution of geographic factors, improves the knowledge on connectivity by considering both short- and long-distance migrants, and provides empirical evidence that migratory connectivity may also depend on migratory dispersion. This has critical implications from a practical point of view because, together with other drivers of migratory connectivity, migratory dispersion has been related to population declines in birds. However, whilst the effect of migratory dispersion is clear, with European migrants occupying larger nonbreeding ranges relative to breeding being less likely to decline (Gilroy et al. 2016, Howard et al. 2020), the effects of migration distance and nonbreeding population spread on avian population dynamics are still obscure. Among European breeding species, migrants with larger nonbreeding population spread seem more likely to have declining populations (Patchett et al. 2018), but confirmation is lacking (Koleček et al. 2018) and an opposite pattern has been shown in the Neotropic migration system (Patchett et al. 2018). Some studies have also suggested more population declines for long-distance European migrants than for shortdistance or resident ones (Sanderson et al. 2006, Vickery et al. 2014, Howard et al. 2020), but others have shown that migration distance does not influence population trends (Gilroy et al. 2016; Patchett et al. 2018) or have suggested that short migration distances reflect a lower adaptive capacity to environmental changes (La Sorte and Fink 2016). Thus, in order to understand how migratory connectivity may impact on bird population dynamics, our study clearly highlights the necessity that migratory connectivity effects should be teased apart from those of migratory dispersion, migration distance and nonbreeding population spread, as each of these players is linked to the others and may have direct, indirect, and potentially divergent consequences on population trends. To this end, the complex interplay between drivers of migratory connectivity unravelled by our analysis must be taken into account to determine how avian population mixing may change in space and time.

## Acknowledgments

We are grateful to A. Alessi for assistance with the INDACO platform, the big data computing facility at the University of Milano. We thank BirdLife International for providing us with bird distribution maps. Special thanks go to all members of the Eurasian African Bird Migration Atlas project team and the European Union for Bird Ringing (EURING), for backing and supporting our research. Funding was provided by the Italian Government (Ministry of the Ecological Transition, formerly Ministry of the Environment, Land and Sea) through a grant to the Convention on Migratory Species (CMS). The authors declare they have no conflicts of interest.

