## Supporting Information for "Eco-evolutionary drivers of avian migratory connectivity"

**Appendix S1.** *Detailed hypotheses and predictions about the determinants of migratory connectivity*

Because migration costs should be minimized when birds relocate between breeding and nonbreeding grounds, we expect that migratory connectivity would decrease with migration distance (Patchett et al. 2018; Finch et al. 2018; Somveille et al. 2021). Geographical constrains related to the configuration of European and African continents also lead us to predict a decrease in migratory connectivity at lower nonbreeding latitude (i.e. more southwards), since populations should be forced to mix more during the nonbreeding period at southern latitudes due to decreasing land availability (Finch et al. 2018). In fact, a large inter-individual spread in the nonbreeding grounds should contribute *per se* to decrease the chance of mixing (Finch et al. 2018). However, the relative configuration of population spreads in the breeding and nonbreeding grounds, also known as 'migratory dispersion', might also shape migratory connectivity; indeed, despite empirical evidence is still lacking, simulation studies suggest that population mixing should increase as the ratio between the latitudinal spreads of the breeding and the nonbreeding grounds departs from equality (Gilroy et al. 2016).

Amongst species-specific traits, low niche specialization is common in long-distance migrants, as it can be advantageous due to the variety of habitats and food resources met *en route* and on the nonbreeding grounds (Reif et al. 2016). If so, natural selection could have maintained a stronger migratory connectivity in specialized species in terms of habitat and dietary breadths, whereas a weaker migratory connectivity should be associated to generalists (Cresswell 2014). Body mass is a fundamental predictor of life-history traits and might affect migratory connectivity in several ways. Larger birds generally show longer migration distances due to an optimization of the aerial locomotion costs (Hein et al. 2012), which may exert an indirect effect on migratory connectivity by increasing population mixing. However, larger species also live longer and, in avian migrants, a longer lifespan promotes the social transmission of migratory routes towards nonbreeding sites and helps maintaining migratory knowledge across generations (Teitelbaum et al. 2016; Foss-Grant et al. 2018). Once the effect of migration distance is accounted for, a larger body mass could thus be expected to favour the evolution of a stronger migratory connectivity. Avian clutch size is a good proxy of *per capita* food availability in the breeding season resulting from the *surplus* of resources allocated for reproduction (Lundblad & Conway 2021). During their life-history, species with larger clutch size should have thus experienced a lower degree of resource competition in the breeding season. As birds relocate in the nonbreeding grounds to minimise the costs arising from resource competition in the breeding grounds (Somveille et al. 2021), a larger clutch size could translate into a stronger migratory connectivity.

Multi-species studies of migratory connectivity have been focused solely on land birds, and comparisons with marine birds are not available. Marine environments are generally associated to less predictable and patchily distributed resources, favouring the evolution of coloniality (Danchin & Wagner 1997). Avian species strictly depending on marine habitats gather in larger colonies compared to terrestrial birds (Møller 2006), with seabirds from larger colonies showing greater migration propensity and longer migration distances because of density-dependent effects of colony size persisting also in the nonbreeding period (Diamond 1978; Lamb et al. 2017). Avoiding intraspecific competition could have promoted relatively higher population mixing in seabirds than in terrestrial birds, thus a lower migratory connectivity may be expected in marine species. Finally, we tested whether migratory connectivity differed between passerine and non-passerine birds. Indeed, the serial residency hypothesis (Cresswell 2014) predicts that birds should tend to redistribute stochastically over a wider nonbreeding area if they are unable to forecast and compensate for any favourable or unfavourable events during migration, particularly juveniles at first migration. We thus expect that passerines would retain a generally lower migratory connectivity than non-passerines because within the former juveniles usually migrate separately from adults and are therefore subject to a larger unpredictability of conditions during migration and in the nonbreeding grounds (Cresswell 2014).

**Appendix S2.** *Selection of ringing encounters*

First, we selected ringing encounter previously validated for their usage in the European-African Bird Migration Atlas (EURING level for 'use.for.atlas' = TRUE). Then, following previous studies on ringing encounters obtained from the EURING Data Bank (e.g. Paradis et al. 1998; Ambrosini et al. 2016), we implemented condition-based criteria in order to reduce encounter heterogeneity. In details, we removed:

1) birds that were not found freshly dead, or birds that were in poor condition or had an accident when ringed, or birds that were alive and probably healthy but taken into captivity (EURING levels for 'condition' = 3, 4, 5, 6);

2) birds that were kept for more than 13 hours during ringing, or birds that have been moved or held extensively during ringing, or those hand reared (EURING levels for 'manipulated' = C, F, T, M, H);

3) birds that were moved unintentionally by man or other agency, or intentionally by man, or moved by water e.g. found on shoreline (EURING levels for 'moved' = 2, 4, 6);

4) birds for which the dates of ringing and/or recovery were not recorded accurately to the nearest 1 week for both the ringing and the finding date (EURING levels for 'date accuracy' = 4, 5, 6, 7, 8);

5) birds for which the places of ringing and/or recovery were not recorded accurately to the nearest 100 km for the ringing or finding places (EURING levels for 'coordinates accuracy' = 6, 7, 8, 9).

Then, we applied a spatiotemporal masking using species-specific criteria in order to remove encounters in non-stationary periods or found within occasional ranges, thus retaining all individuals at their breeding and nonbreeding grounds. Following the phenology reported by Cramp (1998), we identified for each species an "extended" breeding period, corresponding to that spanning the egg breeding for the whole species (even though it may still include *en route* birds for some populations), a "focal" breeding stationary period from the end of spring migration for the latest population to the onset of autumn migration of the earliest population, and a nonbreeding stationary period from the end of autumn migration of the latest population until the onset of spring migration of the earliest population (Table S1). We then removed:

6) encounters found out of the focal breeding and the nonbreeding stationary periods. However, we retained encounters of chicks unable to fledge or of individuals found at nest (EURING levels for 'catching method'

= N and for ‘age by scheme’ = 1) if they occurred during the extended breeding period of each species because we considered that they have occurred in the breeding area of the individual.

Moreover we identified breeding, resident and nonbreeding stationary ranges for each species according to the distribution maps provided by BirdLife International (2019), removing:

7) encounters outside the breeding and resident ranges during the focal or the extended breeding periods;

8) encounters outside the nonbreeding and resident ranges during the nonbreeding stationary period.

After the above step, we checked and manually discarded a few encounters of long-distance migrants remaining outside the European-African migration system, even though within their nonbreeding stationary ranges, to avoid spatial biases in migratory connectivity analysis (e.g. encounters of *Larus* spp. in North America; *Platalea leucorodia* in India; *Sterna hirundo* in Australia and South America). Eventually, we also removed:

9) repeated encounters for the same individual in either the breeding or the nonbreeding range, if any, by retaining the earliest encounter in order to minimise age-bias.

10) individuals that, after the previous steps of data selection, did not have one observation in both the breeding and the nonbreeding ranges.

**Table S1**. Species-specific phenology reported by Cramp (1998): the first integer represents the month, the second one represents the monthly quarter. An exception was made for *Turdus merula*, for which we considered different periods than those reported by Cramp (1998), which were referred mainly to British populations, particularly for the start of the non-breeding period (Santos 1982; Olioso 1995; Main 2002; Andreotti et al. 2010).

| | **Extended breeding period** | | **Focal breeding period** | | **Nonbreeding period** | |
|---|---|---|---|---|---|---|
| **Species** | **Start** | **End** | **Start** | **End** | **Start** | **End** |
| *Anas crecca* | 3-4 | 8-2 | 6-1 | 7-3 | 12-3 | 2-2 |
| *Erithacus rubecula* | 3-1 | 7-3 | 5-2 | 6-4 | 11-3 | 2-3 |
| *Hirundo rustica* | 4-4 | 10-2 | 6-1 | 6-4 | 12-3 | 1-4 |
| *Phalacrocorax carbo* | 2-4 | 10-2 | 4-3 | 6-2 | 12-1 | 1-2 |
| *Cygnus olor* | 4-2 | 11-2 | 4-4 | 8-4 | 1-1 | 2-4 |
| *Larus argentatus* | 4-2 | 8-4 | 5-3 | 8-2 | 12-2 | 2-2 |
| *Larus ridibundus* | 4-1 | 9-1 | 5-2 | 6-2 | 12-1 | 2-2 |
| *Turdus merula* | 2-4 | 9-2 | 5-2 | 7-2 | 12-1 | 2-2 |

| | | | | | | |
|---|---|---|---|---|---|---|
| *Parus caeruleus* | 4-1 | 7-3 | 5-1 | 6-1 | 12-1 | 1-4 |
| *Parus major* | 3-2 | 7-4 | 5-1 | 6-1 | 12-1 | 1-4 |
| *Anas platyrhynchos* | 2-1 | 11-4 | 5-3 | 7-4 | 12-1 | 1-4 |
| *Anser albifrons* | 6-2 | 9-1 | 6-3 | 8-3 | 1-1 | 2-3 |
| *Anser anser* | 3-4 | 8-1 | 4-3 | 7-3 | 12-3 | 1-3 |
| *Accipiter gentilis* | 4-1 | 8-2 | 6-1 | 7-3 | 12-3 | 2-2 |
| *Calidris alpina* | 5-4 | 8-4 | 6-2 | 7-1 | 11-3 | 3-1 |
| *Carduelis chloris* | 4-4 | 7-2 | 5-3 | 7-1 | 12-1 | 2-4 |
| *Carduelis spinus* | 4-3 | 8-3 | 5-3 | 7-4 | 12-3 | 2-2 |
| *Haematopus ostralegus* | 3-4 | 9-3 | 5-1 | 6-2 | 11-1 | 1-4 |
| *Larus canus* | 5-3 | 9-1 | 6-1 | 7-2 | 12-2 | 2-2 |
| *Larus fuscus* | 4-4 | 9-1 | 6-1 | 6-4 | 12-2 | 1-4 |
| *Larus marinus* | 4-3 | 8-4 | 5-2 | 8-2 | 12-2 | 2-2 |
| *Ciconia ciconia* | 4-2 | 8-2 | 5-3 | 7-4 | 12-1 | 1-4 |
| *Cygnus cygnus* | 5-3 | 9-4 | 6-3 | 9-2 | 1-1 | 2-3 |
| *Falco tinnunculus* | 3-4 | 8-2 | 6-1 | 7-3 | 12-3 | 2-2 |
| *Haliaeetus albicilla* | 4-2 | 9-2 | 6-1 | 8-4 | 12-3 | 2-1 |
| *Larus melanocephalus* | 5-2 | 8-3 | 6-1 | 6-4 | 12-1 | 2-4 |
| *Sturnus vulgaris* | 4-2 | 6-2 | 5-1 | 6-1 | 12-1 | 1-4 |
| *Turdus philomelos* | 2-4 | 9-1 | 5-3 | 8-2 | 11-2 | 2-3 |
| *Phalacrocorax aristotelis* | 2-3 | 10-1 | 4-3 | 6-4 | 12-1 | 1-1 |
| *Platalea leucorodia* | 4-1 | 9-2 | 6-1 | 7-3 | 11-4 | 1-4 |
| *Riparia riparia* | 4-4 | 8-4 | 6-1 | 6-4 | 12-1 | 2-3 |
| *Sylvia atricapilla* | 4-3 | 8-1 | 6-3 | 7-2 | 12-1 | 1-3 |
| *Tyto alba* | 2-4 | 12-1 | 4-1 | 10-4 | 12-2 | 2-3 |
| *Alauda arvensis* | 3-3 | 8-4 | 5-1 | 7-4 | 12-1 | 1-3 |
| *Anas acuta* | 4-1 | 8-3 | 6-1 | 7-4 | 12-3 | 1-4 |
| *Anser brachyrhynchus* | 5-2 | 8-4 | 5-3 | 8-2 | 12-1 | 3-2 |
| *Anser fabalis* | 5-3 | 9-1 | 6-1 | 8-3 | 1-1 | 2-4 |
| *Aythya ferina* | 4-3 | 8-2 | 5-3 | 7-4 | 12-2 | 1-4 |
| *Aythya fuligula* | 5-2 | 9-3 | 5-3 | 8-4 | 1-1 | 2-3 |
| *Aythya marila* | 5-3 | 9-3 | 5-4 | 8-2 | 12-1 | 2-3 |
| *Branta canadensis* | 3-3 | 6-4 | 4-1 | 7-2 | 12-1 | 1-4 |
| *Bucephala clangula* | 5-2 | 9-2 | 5-3 | 8-3 | 1-1 | 2-2 |
| *Columba oenas* | 4-3 | 10-4 | 5-1 | 8-2 | 12-1 | 1-4 |
| *Columba palumbus* | 2-3 | 12-1 | 5-2 | 8-3 | 12-2 | 2-2 |
| *Corvus corone* | 4-1 | 6-2 | 5-1 | 5-4 | 11-4 | 2-2 |
| *Corvus frugilegus* | 3-1 | 5-3 | 4-2 | 5-1 | 12-1 | 2-2 |
| *Corvus monedula* | 4-2 | 6-3 | 4-4 | 6-1 | 11-3 | 2-2 |
| *Fulica atra* | 2-4 | 10-1 | 5-3 | 8-2 | 12-2 | 2-3 |
| *Gallinago gallinago* | 3-4 | 9-3 | 5-1 | 7-2 | 12-1 | 1-4 |
| *Gallinula chloropus* | 4-3 | 8-3 | 5-3 | 7-2 | 12-3 | 2-3 |
| *Garrulus glandarius* | 3-4 | 7-4 | 6-2 | 6-4 | 11-2 | 3-3 |
| *Limosa limosa* | 4-1 | 8-3 | 5-2 | 6-3 | 11-1 | 2-1 |
| *Anas penelope* | 5-3 | 8-4 | 6-1 | 8-2 | 12-3 | 2-4 |
| *Anas strepera* | 4-3 | 8-3 | 5-3 | 7-4 | 12-3 | 2-4 |
| *Mergus merganser* | 5-2 | 9-3 | 6-2 | 9-1 | 1-1 | 2-4 |

| | | | | | | |
|---|---|---|---|---|---|---|
| *Netta rufina* | 4-4 | 8-3 | 5-3 | 8-1 | 12-2 | 2-2 |
| *Numenius arquata* | 4-4 | 9-1 | 5-2 | 6-4 | 11-3 | 2-3 |
| *Pluvialis apricaria* | 5-2 | 9-3 | 6-3 | 7-4 | 1-1 | 2-2 |
| *Scolopax rusticola* | 3-2 | 10-3 | 5-2 | 8-3 | 12-1 | 2-3 |
| *Somateria mollissima* | 4-3 | 9-3 | 5-2 | 8-4 | 12-2 | 2-3 |
| *Anas clypeata* | 4-4 | 8-3 | 5-3 | 8-1 | 12-2 | 2-2 |
| *Streptopelia decaocto* | 5-1 | 10-1 | 6-1 | 7-3 | 12-1 | 2-1 |
| *Tetrao urogallus* | 4-3 | 9-2 | 5-3 | 8-2 | 9-3 | 4-2 |
| *Tringa totanus* | 3-4 | 8-3 | 5-1 | 6-2 | 11-1 | 2-2 |
| *Turdus iliacus* | 4-4 | 8-1 | 6-2 | 7-2 | 12-2 | 2-3 |
| *Turdus pilaris* | 4-4 | 8-4 | 5-2 | 8-1 | 12-2 | 2-3 |
| *Turdus viscivorus* | 3-3 | 7-3 | 4-4 | 6-4 | 11-2 | 2-2 |
| *Vanellus vanellus* | 3-1 | 9-2 | 4-2 | 6-2 | 12-2 | 1-3 |
| *Carduelis flammea* | 4-3 | 7-4 | 5-3 | 7-2 | 12-3 | 2-3 |
| *Accipiter nisus* | 4-4 | 8-2 | 6-1 | 7-4 | 12-1 | 2-3 |
| *Acrocephalus melanopogon* | 4-4 | 8-2 | 6-1 | 7-4 | 12-2 | 2-2 |
| *Aegithalos caudatus* | 4-2 | 9-1 | 5-2 | 8-1 | 11-2 | 3-3 |
| *Aegolius funereus* | 2-4 | 8-3 | 5-1 | 7-4 | 12-1 | 2-4 |
| *Aix galericulata* | 4-3 | 8-1 | 4-4 | 7-3 | 8-2 | 4-2 |
| *Alca torda* | 4-3 | 8-2 | 5-2 | 7-2 | 11-1 | 2-2 |
| *Alcedo atthis* | 4-3 | 10-1 | 6-1 | 7-2 | 12-2 | 2-4 |
| *Anthus pratensis* | 3-4 | 8-4 | 5-3 | 6-2 | 12-1 | 1-4 |
| *Aquila chrysaetos* | 3-1 | 8-2 | 4-2 | 7-4 | 12-2 | 2-4 |
| *Ardea cinerea* | 3-3 | 8-2 | 4-1 | 6-2 | 12-1 | 1-2 |
| *Asio otus* | 2-4 | 7-4 | 6-3 | 7-2 | 12-3 | 2-3 |
| *Athene noctua* | 3-3 | 8-3 | 4-2 | 7-4 | 8-4 | 3-2 |
| *Branta leucopsis* | 5-4 | 8-4 | 6-1 | 8-2 | 1-1 | 3-3 |
| *Buteo buteo* | 3-4 | 8-1 | 5-3 | 7-3 | 12-1 | 1-4 |
| *Carduelis carduelis* | 5-1 | 8-4 | 5-4 | 8-1 | 12-2 | 2-4 |
| *Certhia brachydactyla* | 3-4 | 7-4 | 4-3 | 7-2 | 8-1 | 3-3 |
| *Certhia familiaris* | 3-4 | 7-4 | 5-1 | 6-4 | 12-1 | 2-4 |
| *Cettia cetti* | 6-1 | 8-4 | 6-3 | 8-2 | 11-1 | 4-4 |
| *Charadrius alexandrinus* | 4-2 | 9-1 | 5-4 | 6-4 | 11-3 | 2-2 |
| *Charadrius hiaticula* | 3-4 | 9-3 | 5-3 | 7-4 | 12-1 | 1-4 |
| *Cinclus cinclus* | 2-4 | 9-1 | 6-1 | 8-2 | 11-1 | 1-4 |
| *Coccothraustes coccothraustes* | 4-2 | 8-4 | 4-3 | 8-3 | 12-2 | 1-4 |
| *Corvus corax* | 1-4 | 8-4 | 4-1 | 6-3 | 9-1 | 1-3 |
| *Luscinia svecica* | 4-4 | 8-2 | 6-2 | 7-4 | 12-1 | 1-4 |
| *Dendrocopos major* | 4-3 | 7-3 | 5-1 | 7-1 | 11-2 | 2-4 |
| *Emberiza citrinella* | 4-3 | 8-2 | 5-3 | 7-4 | 11-3 | 3-2 |
| *Emberiza schoeniclus* | 5-1 | 7-3 | 5-3 | 7-1 | 11-3 | 2-2 |
| *Falco peregrinus* | 5-1 | 8-3 | 5-3 | 7-4 | 11-3 | 2-3 |
| *Fratercula arctica* | 4-1 | 8-4 | 4-3 | 7-4 | 10-2 | 3-2 |
| *Fringilla coelebs* | 4-3 | 7-2 | 5-4 | 6-3 | 12-1 | 2-3 |
| *Fringilla montifringilla* | 5-2 | 7-3 | 6-2 | 7-1 | 11-3 | 2-3 |
| *Glaucidium passerinum* | 4-3 | 7-4 | 4-4 | 7-2 | 12-2 | 2-4 |
| *Grus grus* | 4-4 | 9-2 | 5-3 | 7-2 | 12-1 | 2-2 |

| | | | | | | |
|---|---|---|---|---|---|---|
| *Sterna caspia* | 5-1 | 8-3 | 5-4 | 7-2 | 12-3 | 2-3 |
| *Larus audouinii* | 4-3 | 8-3 | 5-1 | 7-4 | 11-3 | 2-2 |
| *Carduelis cannabina* | 4-3 | 8-4 | 5-3 | 7-4 | 11-3 | 2-2 |
| *Carduelis flavirostris* | 4-1 | 8-3 | 5-3 | 7-4 | 12-3 | 2-3 |
| *Milvus milvus* | 3-4 | 7-4 | 5-3 | 7-2 | 12-1 | 1-4 |
| *Sula bassana* | 4-2 | 11-1 | 6-2 | 7-4 | 12-1 | 1-4 |
| *Motacilla alba* | 4-1 | 8-2 | 6-1 | 7-3 | 11-3 | 1-4 |
| *Nucifraga caryocatactes* | 2-4 | 7-4 | 4-1 | 5-4 | 11-1 | 2-3 |
| *Pandion haliaetus* | 4-3 | 8-1 | 6-1 | 7-4 | 12-1 | 2-3 |
| *Panurus biarmicus* | 3-3 | 7-1 | 4-3 | 6-3 | 12-1 | 2-3 |
| *Passer hispaniolensis* | 3-1 | 10-4 | 5-3 | 8-3 | 12-1 | 2-4 |
| *Passer montanus* | 4-1 | 9-2 | 4-4 | 8-2 | 11-3 | 3-3 |
| *Parus ater* | 4-2 | 7-4 | 4-4 | 6-3 | 10-3 | 2-4 |
| *Phoenicurus ochruros* | 4-3 | 7-4 | 5-1 | 7-1 | 11-3 | 2-3 |
| *Phylloscopus collybita* | 4-4 | 8-1 | 6-1 | 7-2 | 11-4 | 2-3 |
| *Podiceps cristatus* | 2-3 | 9-1 | 5-3 | 7-3 | 12-3 | 1-4 |
| *Parus montanus* | 4-3 | 8-1 | 5-3 | 6-1 | 12-1 | 3-1 |
| *Prunella modularis* | 3-1 | 9-2 | 6-1 | 8-2 | 11-3 | 2-3 |
| *Pyrrhula pyrrhula* | 4-3 | 8-3 | 5-2 | 7-3 | 12-1 | 2-3 |
| *Recurvirostra avosetta* | 4-2 | 9-1 | 5-3 | 7-2 | 11-2 | 2-2 |
| *Regulus ignicapillus* | 4-2 | 8-2 | 5-3 | 7-4 | 12-3 | 1-4 |
| *Regulus regulus* | 4-3 | 8-3 | 6-1 | 7-3 | 12-1 | 2-3 |
| *Remiz pendulinus* | 4-4 | 8-3 | 5-1 | 6-4 | 12-3 | 1-4 |
| *Rissa tridactyla* | 5-2 | 9-2 | 6-1 | 7-4 | 12-1 | 1-4 |
| *Saxicola torquata* | 4-2 | 8-2 | 6-1 | 7-4 | 11-4 | 1-3 |
| *Serinus serinus* | 4-1 | 8-2 | 5-1 | 7-3 | 11-2 | 2-3 |
| *Sitta europaea* | 3-4 | 7-3 | 5-1 | 6-2 | 11-1 | 3-3 |
| *Sterna hirundo* | 5-2 | 9-2 | 6-3 | 7-2 | 11-2 | 3-1 |
| *Sterna albifrons* | 5-2 | 9-1 | 6-1 | 7-2 | 11-2 | 3-1 |
| *Strix aluco* | 2-3 | 7-1 | 2-4 | 6-3 | 7-4 | 2-2 |
| *Sylvia melanocephala* | 3-3 | 7-2 | 5-1 | 5-4 | 1-1 | 2-2 |
| *Tadorna tadorna* | 4-3 | 9-1 | 5-1 | 8-3 | 1-1 | 2-2 |
| *Sterna sandvicensis* | 4-4 | 7-4 | 5-2 | 6-4 | 12-1 | 1-2 |
| *Troglodytes troglodytes* | 3-4 | 8-3 | 5-4 | 7-4 | 12-3 | 3-2 |
| *Uria aalge* | 4-4 | 8-1 | 5-1 | 7-2 | 12-1 | 1-4 |

**Appendix S3.** *Discrimination of migratory and resident geographical populations*

The distinction between migratory and non-migratory species or populations is particularly challenging for birds, where different types of migration are exhibited and large differences exist, even between populations of the same species (Chapman et al. 2011). In this work, we avoided to classify migratory *vs* resident geographical populations (i.e. clusters identified by the migratory connectivity analysis) on the basis of a cut-off migration distance, because many populations of birds are known to migrate even when migration distance is short, and migration distance is often population-specific as well as is influenced by various ecological and climatic drivers (Visser et al. 2009; Meller et al. 2016; Curley et al. 2020). Indeed, migration distance is only one amongst other many characteristics that are used to define migration, such as periodicity or directionality of movements (Eyres et al. 2017). Consequently, we implemented a classification based on the inspection of the overall spatial pattern of individual positions observed in the non-breeding stationary range, (i.e. after migration), relative to that found in the breeding range, i.e. prior to migration. For each geographical population, we first calculated the 95% minimum convex polygon (MCP) of individual locations in the breeding period (breeding MCP) and the 95% MCP of the same individual locations in the non-breeding stationary period (non-breeding MCP). We then overlapped the breeding and non-breeding MCPs and classified those where the overlap was more than 75% of the area of the breeding MCP as resident populations. Populations were classified as migratory otherwise. Such classification should be helpful especially to distinguish sedentary, or non-migratory geographical populations with a general tendency of dispersal, from those showing actual migration patterns. MCP has been used to quantify population spread of birds in the breeding and non-breeding grounds (e.g. Blackburn et al. 2017; Burgess et al. 2020) and we should note that, in our case, the increase in MCP extent with increasing sample size (Burgess et al. 2020) would not affect the classification outcome because both the MCPs include exactly the same number of individuals.

**Appendix S4.** *Species investigated in phylogenetic comparative meta-analysis*

**Table S2**. Number of geographical populations (i.e., clusters) and individuals for each bird species included in the phylogenetic comparative meta-analysis of the strength of migratory connectivity.

| Family | Species | N geographical populations | N individuals |
|---|---|---|---|
| Accipitridae | *Accipiter nisus* | 6 | 3481 |
| | *Aquila chrysaetos* | 2 | 437 |
| | *Buteo buteo* | 1 | 3509 |
| | *Haliaeetus albicilla* | 2 | 2629 |
| | *Milvus milvus* | 2 | 862 |
| Alaudidae | *Alauda arvensis* | 1 | 53 |
| Alcidae | *Alca torda* | 2 | 521 |
| | *Fratercula arctica* | 2 | 193 |
| Anatidae | *Anas acuta* | 1 | 34 |
| | *Anas clypeata* | 1 | 278 |
| | *Anas crecca* | 1 | 305 |
| | *Anas penelope* | 3 | 147 |
| | *Anas platyrhynchos* | 1 | 6333 |
| | *Anas strepera* | 1 | 234 |
| | *Anser anser* | 3 | 157 |
| | *Anser brachyrhynchus* | 2 | 1089 |
| | *Anser fabalis* | 2 | 289 |
| | *Aythya ferina* | 1 | 552 |
| | *Aythya fuligula* | 2 | 1095 |
| | *Branta leucopsis* | 3 | 402 |
| | *Bucephala clangula* | 1 | 69 |
| | *Cygnus cygnus* | 2 | 529 |
| | *Somateria mollissima* | 3 | 1182 |
| | *Tadorna tadorna* | 5 | 310 |
| Charadriidae | *Charadrius hiaticula* | 1 | 42 |
| | *Vanellus vanellus* | 1 | 3646 |
| Ciconiidae | *Ciconia ciconia* | 2 | 3783 |
| Columbidae | *Columba oenas* | 2 | 345 |
| | *Columba palumbus* | 1 | 1522 |
| Corvidae | *Corvus corone* | 1 | 1264 |
| | *Corvus frugilegus* | 1 | 517 |
| | *Corvus monedula* | 9 | 1557 |
| Emberizidae | *Emberiza schoeniclus* | 2 | 937 |
| Falconidae | *Falco peregrinus* | 3 | 201 |

| | | | |
|---|---|---|---|
| | *Falco tinnunculus* | 5 | 7662 |
| Fringillidae | *Carduelis cannabina* | 1 | 491 |
| | *Carduelis flammea* | 2 | 202 |
| | *Carduelis flavirostris* | 2 | 66 |
| | *Carduelis spinus* | 2 | 1370 |
| | *Coccothraustes coccothraustes* | 7 | 571 |
| | *Fringilla coelebs* | 1 | 2682 |
| | *Serinus serinus* | 2 | 672 |
| Gruidae | *Grus grus* | 1 | 372 |
| Haematopodidae | *Haematopus ostralegus* | 1 | 2349 |
| Hirundinidae | *Hirundo rustica* | 2 | 96 |
| Laridae | *Larus argentatus* | 1 | 19,823 |
| | *Larus canus* | 1 | 3878 |
| | *Larus fuscus* | 1 | 5015 |
| | *Larus marinus* | 6 | 2380 |
| | *Larus melanocephalus* | 9 | 1099 |
| | *Larus ridibundus* | 1 | 27,479 |
| | *Rissa tridactyla* | 1 | 134 |
| Motacillidae | *Anthus pratensis* | 1 | 51 |
| | *Motacilla alba* | 2 | 439 |
| Muscicapidae | *Erithacus rubecula* | 2 | 6621 |
| | *Luscinia svecica* | 1 | 35 |
| | *Phoenicurus ochruros* | 2 | 306 |
| | *Saxicola torquata* | 2 | 65 |
| Pandionidae | *Pandion haliaetus* | 1 | 369 |
| Paridae | *Panurus biarmicus* | 2 | 145 |
| Phalacrocoracidae | *Phalacrocorax aristotelis* | 4 | 916 |
| | *Phalacrocorax carbo* | 1 | 2033 |
| Phylloscopidae | *Phylloscopus collybita* | 1 | 133 |
| Rallidae | *Fulica atra* | 9 | 3806 |
| | *Gallinula chloropus* | 2 | 528 |
| Recurvirostridae | *Recurvirostra avosetta* | 2 | 372 |
| Remizidae | *Remiz pendulinus* | 1 | 53 |
| Scolopacidae | *Gallinago gallinago* | 1 | 137 |
| | *Limosa limosa* | 2 | 165 |
| | *Numenius arquata* | 2 | 485 |
| | *Scolopax rusticola* | 1 | 518 |
| | *Tringa totanus* | 4 | 363 |
| Sternidae | *Sterna hirundo* | 2 | 1089 |
| | *Sterna sandvicensis* | 3 | 423 |

| | | | |
|---|---|---|---|
| Strigidae | *Asio otus* | 3 | 618 |
| Sturnidae | *Sturnus vulgaris* | 3 | 7038 |
| Sulidae | *Sula bassana* | 1 | 350 |
| Sylviidae | *Sylvia atricapilla* | 4 | 255 |
| Threskiornithidae | *Platalea leucorodia* | 7 | 817 |
| Turdidae | *Turdus iliacus* | 1 | 683 |
| | *Turdus philomelos* | 3 | 5137 |
| | *Turdus pilaris* | 1 | 1572 |
| | *Turdus viscivorus* | 1 | 542 |

**Appendix S5.** *Compilation of species-specific life-history traits*

Species-specific life-history traits entered as fixed effects in the meta-analysis were: diet breadth, habitat breadth, whether a specie was marine or terrestrial, body mass, and clutch size.

Information on diet breadth was obtained from del Hoyo et al. (2017). We considered the following food sources: 1) terrestrial invertebrates, 2) aquatic invertebrates, 3) terrestrial vertebrates, 4) aquatic vertebrates, 5) fruits and berries, 6) sedges and seeds, 7) other plant materials (e.g. nectar, grass or aquatic plants) and counted the number of them representing a substantial contribution to a species diet to obtain an estimate of the diet breadth.

Similarly, habitat breadth was estimated as the total number of primary habitat types exploited by a species, as reported in BirdLife (2020). Species were also considered as marine when BirdLife (2020), reported that ‘marine habitat’ was their primary habitat and non-marine otherwise. Body mass and clutch size were also compiled from BirdLife (2020).

**Appendix S6.** *Moderators entered in the phylogenetic meta-analytic model*

**Table S3**. Information on moderators initially considered as fixed effects for the phylogenetic meta-analytic mixed model concerning 191 geographical populations of 83 bird species. For relevant hypotheses, see Table 1 in the main text.

| **Moderator (units)** | **Type** | **Transformation** | **Level** | **Notes** |
|---|---|---|---|---|
| Mean migration distance (km) | Continuous | ln(ln(*x*) + 1) | Geographical population | Not collinear with nonbreeding latitude (r = -0.31) or population spread (r = 0.54) |
| Mean nonbreeding latitude (°) | Continuous | – | Geographical population | Not collinear with migration distance (r = -0.31) or population spread (r = -0.35) |
| Mean nonbreeding population spread (km) | Continuous | ln(*x*) | Geographical population | Not collinear with migration distance (r = 0.54) or nonbreeding latitude (r = -0.35) |
| Breeding to nonbreeding ratio in population spread | Continuous | – | Geographical population | – |
| Body mass (grams) | Continuous | – | Species | – |
| Habitat breadth | Integer | – | Species | Not collinear with diet breadth (r = 0.14) |
| Diet breadth | Integer | – | Species | Not collinear with habitat breadth (r = 0.14) |
| Clutch size | Integer | – | Species | – |
| Marine or non-marine | Binary | – | Species | – |
| Passerine or non-passerine | Binary | – | Species | – |

**Appendix S7.** *Preliminary data exploration and model performance*

Visual inspection of the data showed that $Z_r$ values varied non-linearly with mean migration distance. In contrast, the relationship was linear after double-natural logarithm transformation (Fig. S1). The variance of $Z_r$ values also markedly differed between marine and non-marine species and between passerine and non-passerines (Fig. S2). $Z_r$ values predicted from the full model showed a good agreement with observed ones indicating a proper fit of the model (Fig. S3).

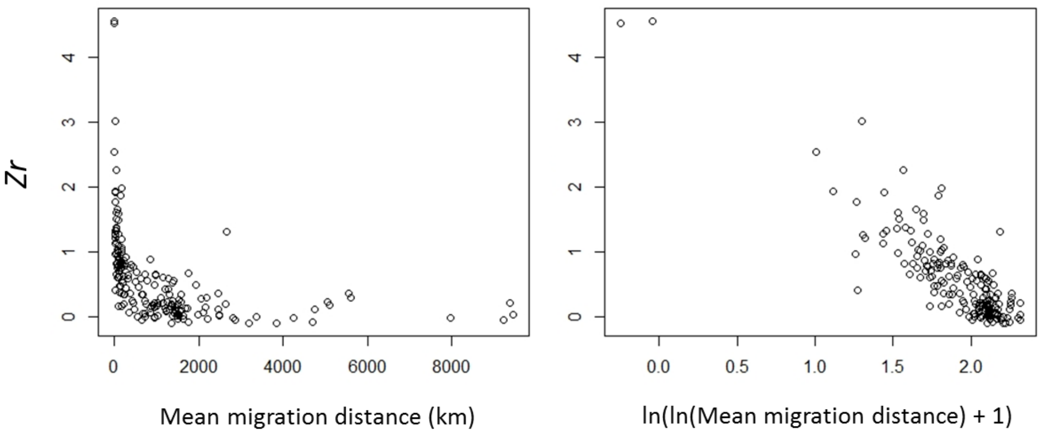


**Figure S1**. $Z_r$ values according to mean migration distance before and after the double logarithmic transformation. Note that a constant term (1) was added to avoid negative values in the second logarithm.

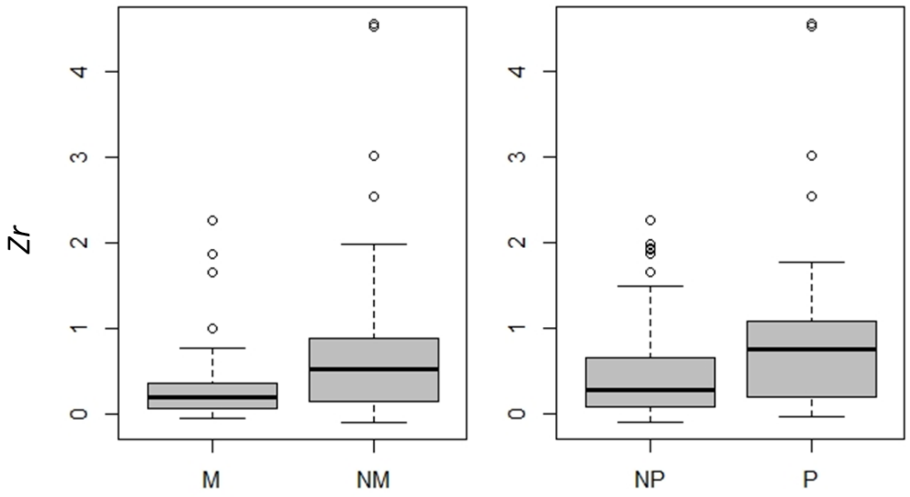


**Figure S2**. $Z_r$ values of marine (M) and non-marine (NM) species and of passerine (P) and non-passerine (NP) birds showing variance heterogeneity between groups.

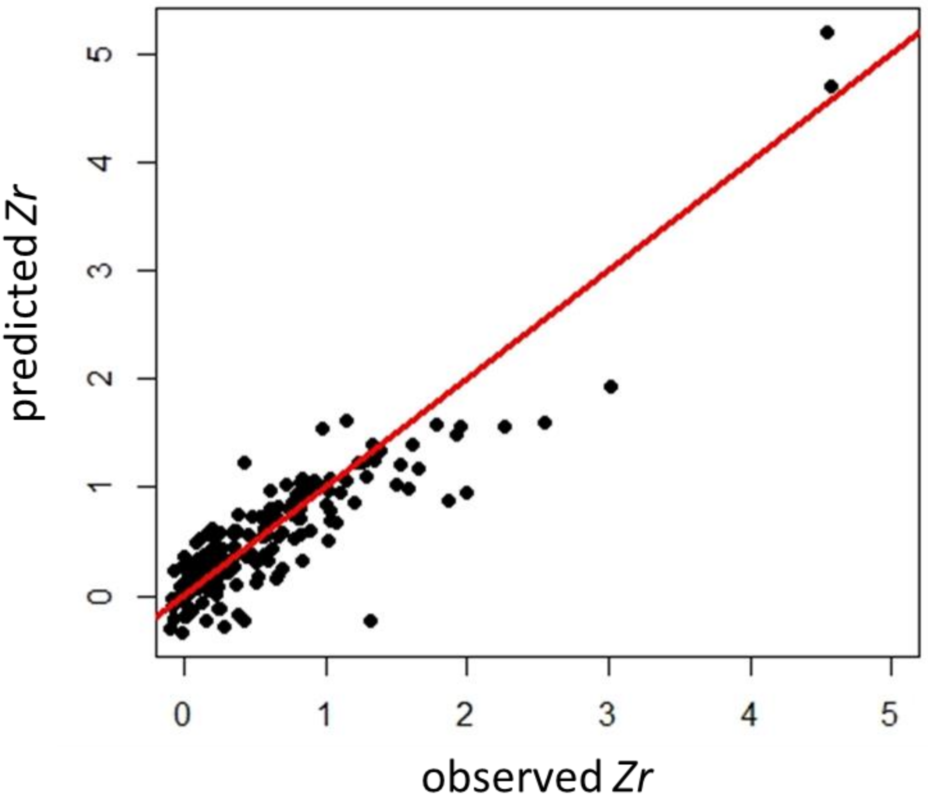


**Figure S3**. Model performance evaluated through inspection of predicted vs. observed $Z_r$ values ($R^2$ = 0.80). Diagonal line is the 1 to 1 line.

**Appendix S8.** *Analysis considering only migratory populations*

Our phylogenetic meta-analytic mixed models were re-run by considering only the geographical populations classified as migratory. For the null model, the results confirmed that migratory connectivity was moderate and significantly larger than zero (estimated *Zr* = 0.397 ± 0.083 SE, $t_{82}$ = 4.769, P < 0.001, corresponding to $r_M$ = 0.377, 95% CI: 0.228 – 0.509; Fig. 2), although there was no significant phylogenetic signal in the strength of migratory connectivity ($H^2$ = 0.146, $\chi^2$ = 2.543, df = 1, P = 0.111).

For the full model, results confirmed the negative effect of migration distance, the positive effect of population spread and the negative effect of the mean nonbreeding latitude on the strength of migratory connectivity (Table S4), whereas the quadratic effect of the ratio between the breeding and nonbreeding population spread and the effect of habitat breadth were not significant (Table S4). There was no phylogenetic signal on migratory connectivity ($H^2$ = 1.596 × $10^{-8}$, $\chi^2$ = 0.000, df = 1, P = 1.000). The model showed a good performance (observed *vs*. predicted values: $R^2$ = 0.67) and the diagnostics indicated that no relevant deviation from assumptions occurred (Appendix S9).

**Table S4**. Parameters estimated from the phylogenetic meta-analytic mixed model explaining the strength of migratory connectivity (as Fisher Z-transformation of $r_M$ value) across 150 migrant geographical populations belonging to 83 species. Transformations were applied to 'Migration distance' and 'Nonbreeding population spread' (see Appendix S6-S7), while the second order polynomial term of 'Migratory dispersion' was included to account for quadratic effects (Table 1). All moderators are scaled. An asterisk marks significant (P < 0.05) moderators.

| Moderator | Coefficient | SE | t | df | P | |
|---|---|---|---|---|---|---|
| Intercept | 0.395 | 0.021 | 18.544 | 71 | < 0.001 | * |
| Migration distance (km) | -0.377 | 0.014 | -27.155 | 138 | < 0.001 | * |
| Nonbreeding population spread (km) | 0.044 | 0.017 | 2.552 | 138 | 0.011 | * |
| Migratory dispersion | -0.006 | 0.015 | -0.417 | 138 | 0.677 | |
| Migratory dispersion$^2$ | <0.001 | 0.003 | 0.134 | 138 | 0.894 | |
| Nonbreeding latitude (°) | -0.039 | 0.012 | -3.275 | 138 | 0.001 | * |
| Body mass (kg) | 0.001 | 0.023 | 0.058 | 71 | 0.954 | |
| Habitat diversity | 0.046 | 0.026 | 1.777 | 71 | 0.078 | |
| Diet diversity | 0.019 | 0.020 | 0.962 | 71 | 0.338 | |
| Clutch size | 0.041 | 0.029 | 1.436 | 71 | 0.153 | |
| Marine | -0.027 | 0.027 | -1.001 | 71 | 0.318 | |
| Passerine | -0.014 | 0.025 | -0.566 | 71 | 0.572 | |

**Appendix S9.** *Model diagnostics*

Even if this study does not represent a proper meta-analysis as information were not retrieved from the literature, we applied a set of techniques typical of meta-analyses to assess the robustness of the results. Indeed, these data are, essentially, based on ringing data, and it is well known that some species or some geographical populations may be under or overrepresented due to the large temporal and spatial variability in encounter probability. We argued that these processes may generate biases in the data similar to those deriving from publication bias in typical meta-analyses and we thus assessed the robustness of the results. Rosenthal's fail-safe number was always very large in both meta-analyses (all geographical populations: 1,787,814; $P < 0.001$; migratory populations only: 561,658; $P < 0.001$). In addition, funnel plots were rather symmetric (Rank Correlation Test for Funnel Plot Asymmetry; all geographical populations: Kendall's tau = 0.0614, P = 0.207; migratory populations only: Kendall's tau = 0.085, P = 0.123; Fig. S4), and the intercepts of Egger's regressions were not significant (all geographical populations: $t_{189} = -0.051$, $P = 0.960$; migratory populations only: $t_{148} = 0.711$, $P = 0.478$). These diagnostics thus indicate that the results of our analyses are robust.

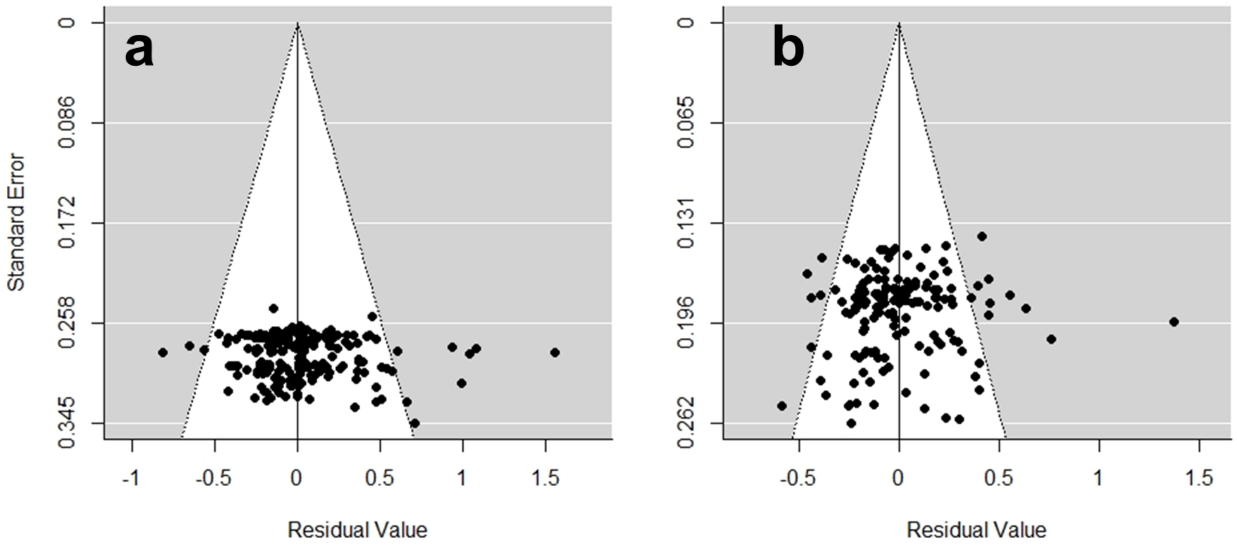


**Figure S4**. Funnel plots for the analyses conducted on all geographical populations (a) and restricted to migrant populations (b).

**Supporting Information - References**


Ambrosini, R., Cuervo, J. J., du Feu, C., Fiedler, W., Musitelli, F., Rubolini, D. et al. (2016). Migratory connectivity and effects of winter temperatures on migratory behaviour of the European robin *Erithacus rubecula*: a continent-wide analysis. *Journal of Animal Ecology*, 85, 749-760.

Andreotti, A., Pirrello, S., Tomasini, S., Merli, F. (2010). I tordi in Italia. *ISPRA Rapporti 123/2010*, 1-153.

BirdLife International (2020). *IUCN Red List for birds*. Downloaded from http://www.birdlife.org on 02/11/2020.

BirdLife International and Handbook of the Birds of the World (2019). *Bird species distribution maps of the world*. Version 2019.1. Available at http://datazone.birdlife.org/species/requestdis.

Blackburn, E., Burgess, M., Freeman, B., Risely, A., Izang, A., Ivande, S., et al. (2017). Low and annually variable migratory connectivity in a long-distance migrant: Whinchats *Saxicola rubetra* may show a bet-hedging strategy. *Ibis*, 159, 902-918.

Burgess, M.D., Finch, T., Border, J.A., Castello, J., Conway, G., Ketcher, M., et al. (2020). Weak migratory connectivity, loop migration and multiple non-breeding site use in British breeding Whinchats *Saxicola rubetra*. *Ibis*, 162, 1292-1302.

Chapman, B.B., Brönmark, C., Nilsson, J.Å., Hansson, L.A. (2011). The ecology and evolution of partial migration. *Oikos*, 120, 1764-1775.

Cramp, S. (1998). *The complete birds of the Western Palearctic on CD-ROM*. Oxford University Press, Oxford.

Cresswell, W. (2014). Migratory connectivity of Palaearctic–African migratory birds and their responses to environmental change: the serial residency hypothesis. *Ibis*, 156, 493-510.

Curley, S.R., Manne, L.L., Veit, R.R. (2020). Differential winter and breeding range shifts: implications for avian migration distances. *Diversity and Distributions*, 26, 415-425.

Danchin, E., Wagner, R.H. (1997). The evolution of coloniality: the emergence of new perspectives. *Trends in Ecology & Evolution*, 12, 342-347.

Del Hoyo, J., Elliott, A., Sargatal, J., Christie, D.A., de Juana, E. (2017). *Handbook of the birds of the world alive*. Lynx Edicions, Barcelona.

Diamond, A.W. (1978). Feeding strategies and population size in tropical seabirds. *The American Naturalist*, 112, 215-223.

Eyres, A., Böhning-Gaese, K., Fritz, S.A. (2017). Quantification of climatic niches in birds: adding the temporal dimension. *Journal of Avian Biology*, 48, 1517-1531.

Finch, T., Butler, S.J., Franco, A.M., Cresswell, W. (2017). Low migratory connectivity is common in long-distance migrant birds. *Journal of Animal Ecology*, 86, 662-673.

Foss-Grant, A., Bewick, S., Fagan, W.F. (2018). Social transmission of migratory knowledge: quantifying the risk of losing migratory behavior. *Theoretical Ecology*, 11, 257-270.

Gilroy, J.J., Gill, J.A., Butchart, S.H., Jones, V.R., Franco, A.M. (2016). Migratory diversity predicts population declines in birds. *Ecology Letters*, 19, 308-317.

Hein, A.M., Hou, C., Gillooly, J.F. (2012). Energetic and biomechanical constraints on animal migration distance. *Ecology Letters*, 15, 104-110.

Lamb, J.S., Satgé, Y.G., Jodice, P.G. (2017). Influence of density-dependent competition on foraging and migratory behavior of a subtropical colonial seabird. *Ecology and Evolution*, 7, 6469-6481.

Lundblad, C.G., Conway, C. J. (2021). Ashmole's hypothesis and the latitudinal gradient in clutch size. *Biological Reviews*, 96, 1349–1366.

Main, I. (2002). Seasonal movements of Fennoscandian Blackbirds *Turdus merula*. *Ringing and Migration*, 21, 65-74.

Meller, K., Vähätalo, A.V., Hokkanen, T., Rintala, J., Piha, M., Lehikoinen, A. (2016). Interannual variation and long-term trends in proportions of resident individuals in partially migratory birds. *Journal of Animal Ecology*, 85, 570-580.

Møller, A.P. (2006). Sociality, age at first reproduction and senescence: comparative analyses of birds. *Journal of Evolutionary Biology*, 19, 682-689.

Olioso, G. (1995). La migration prenutiale des especes du genre Turdus en Provence. Analyse des repreises de bagues. *Fauna de Provence (C.E.E.P.)*, 16, 73-85.

Paradis, E., Baillie, S.R., Sutherland, W.J., Gregory, R.D. (1998). Patterns of natal and breeding dispersal in birds. *Journal of Animal Ecology*, 67, 518–536.

Patchett, R., Finch, T., Cresswell, W. (2018). Population consequences of migratory variability differ between flyways. *Current Biology*, 28, R340-R341.

Reif, J., Hořák, D., Krištín, A., Kopsová, L., Devictor, V. (2016). Linking habitat specialization with species' traits in European birds. *Oikos*, 125, 405-413.

Santos, T. (1982) *Migracion e invernada de zorzales y mirlos (genero Turdus) en la Peninsula Iberica*. Tesi doctoral. Ed. de la Universidad Complutense, Madrid, Spain.

Somveille, M., Bay, R.A., Smith, T.B., Marra, P.P., Ruegg, K.C. (2021). A general theory of avian migratory connectivity. *Ecology Letters*, 24, 1848-1858.

Teitelbaum, C.S., Converse, S.J., Fagan, W.F., Böhning-Gaese, K., O'Hara, R.B., Lacy, A.E., Mueller, T. (2016). Experience drives innovation of new migration patterns of whooping cranes in response to global change. *Nature Communications*, 7, 1-7.

Visser, M.E., Perdeck, A.C., Van Balen, J.H., Both, C. (2009). Climate change leads to decreasing bird migration distances. *Global Change Biology*, 15, 1859-1865.
